# Leukemia-derived apelin selects endothelial niche clones to promote tumorigenesis

**DOI:** 10.1101/2024.09.09.612077

**Authors:** Chloé S. Baron, Olivia Mitchell, Serine Avagyan, Romain Menard, Song Yang, Anne L. Robertson, Rajiv Potluri, Jay Shendure, Romain Madelaine, Aaron McKenna, Leonard I. Zon

## Abstract

Hematopoietic stem cells are regulated by endothelial and mesenchymal stromal cells in the marrow niche^1–3^. Leukemogenesis was long believed to be solely driven by genetic perturbations in hematopoietic cells but introduction of genetic mutations in the microenvironment demonstrated the ability of niche cells to drive disease progression^4–8^. The mechanisms by which the stem cell niche induces leukemia remain poorly understood. Here, using cellular barcoding in zebrafish, we found that clones of niche endothelial and stromal cells are significantly expanded in leukemic marrows. The pro-angiogenic peptide *apelin* secreted by leukemic cells induced sinusoidal endothelial cell clonal selection and transcriptional reprogramming towards an angiogenic state to promote leukemogenesis *in vivo*. Overexpression of *apelin* in normal hematopoietic stem cells led to clonal amplification of the niche endothelial cells and promotes clonal dominance of blood cells. Knock-out of *apelin* in leukemic zebrafish resulted in a significant reduction in disease progression. Our results demonstrate that leukemic cells remodel the clonal and transcriptional landscape of the marrow niche to promote leukemogenesis and provide a potential therapeutic opportunity for anti-*apelin* treatment.

## Main text

Hematopoietic stem cells (HSCs) are essential to produce all mature blood cells during the entire life of an organism^9,10^. HSCs reside in a complex microenvironment or niche that provides signals for their differentiation and self-renewal^11–13^. Endothelial and mesenchymal stromal cells (ECs and MSCs, respectively) are two key cellular components of the stem cell niche that are indispensable for HSC function^1–3,14^. Most HSCs reside near sinusoidal ECs and ablation of ECs results in HSC loss highlighting the crucial role of ECs in the niche^15,16^. A rare subset of niche ECs expresses *apelin*, an angiogenic growth factor necessary for HSC function at steady state^17,18^. *Apelin*^+^ ECs possess endothelial progenitor potential that promotes vascular regeneration after irradiation-induced damage and is necessary for HSC engraftment upon transplantation^18^. MSCs are a cell type tightly associated with HSC function as specific ablation of MSC secreting high levels of CXCL12 was demonstrated to deplete HSCs from the niche^19,20^. Endothelial and stromal cells provide an environment that is conducive to the growth and function of normal HSCs.

Malignant transformation and abnormal proliferation of an HSC clone results in hematological malignancies such as acute myeloid leukemia (AML)^21^. The HSC niche has been proposed to contribute to leukemogenesis as genetic alterations of APC, Dicer or RB in niche cells have been shown to induce myelodysplastic syndrome (MDS) and pre-leukemic states^4–8^. Leukemia induces extensive niche remodeling including angiogenesis, a process by which new blood vessels are formed. At steady state, angiogenesis is tightly regulated by a balance of pro- and anti-angiogenic factors. Active and abnormal angiogenesis has been demonstrated in the marrow of patients with diverse subtypes of acute and chronic leukemia^22–26^. Studies showed that leukemic cells secrete pro-angiogenic factors that stimulate ECs to support leukemic growth^24^. In this study, we used a CRISPR-Cas9 based genetic lineage tracing method^27,28^ paired with single-cell mRNA-Sequencing (scRNA-seq) to investigate the clonal and transcriptional response of niche ECs and MSCs upon leukemia formation *in vivo*. Using genetic overexpression and knock-out zebrafish models and human primary samples, we demonstrate that *apelin* secreted by leukemic cells induces selection of niche endothelial clones and transcriptional remodeling towards an endothelial progenitor state driving abnormal angiogenesis. We link this endothelial clonal selection via *apelin* signaling to leukemia progression demonstrating for the first time that excessive production of a secreted ligand by leukemic cells remodels the endothelial niche to support leukemogenesis.

### Overexpression of human *MYC* in blood cells leads to acute myeloid leukemia in zebrafish

To generate a myeloid leukemia model in zebrafish, we mosaically overexpressed the human oncogene *MYC* under the early blood lineage specific draculin promotor (*drl:MYC*)^29^. D*rl:MYC* injected zebrafish died significantly quicker than non-injected controls (**Fig. 1A**). These animals were smaller in size, exhibiting bleeding, cardiac oedema secondary to anemia and difficulty swimming (**Fig. 1B**). Flow cytometry analysis demonstrated that *drl:MYC* marrows had leukemic blast-like population that overlapped with the progenitor gate in healthy marrows and a significant decrease of mature myeloid, lymphoid and erythroid cells (as defined in forward and side scatter space, FSC and SSC, respectively^30^, **Fig. 1C**; **Fig. S1A**). Large pro-erythroblasts were sometimes detected instead of or in combination with leukemic cells (**Fig. S1A**). Staining of total marrows with May-Grünwald Giemsa solution revealed abundant dark purple leukemic cells with a high nuclear to cytoplasmic ratio compared to control (**Fig. 1D**). Flow cytometry analysis of peripheral blood samples also detected circulating blasts (**Fig. S1B**). Immunohistochemistry on whole marrow sections demonstrated a high abundance of MYC^+^ leukemic cells (**Fig. S1C**). Bulk RNA-Sequencing of control *drl:mCherry* and *drl:MYC-mCherry* total marrow cells revealed a significant upregulation of embryonic hemoglobins *hbbe1.1* and *hbbe1.2* and the master erythroid transcription factor *gata1a* confirming the erythroid identity of leukemic cells (**Fig. 1E**). The embryonic globin expression suggested the cell of origin was developmental in nature. We validated this result using the zebrafish transgenic lines *lcr:GFP* and *gata1a:dsRed* which label mature erythrocytes and erythroid progenitors, respectively, and demonstrated that *drl:MYC* marrows are significantly enriched in *lcr:GFP*^+^;*gata1:dsRed*^+^ leukemic cells and/or *lcr:GFP*^+^;*gata1:dsRed*^-^ pro-erythroblasts revealing leukemic transformation at different stages of erythropoiesis (**Fig. S1D**)^31,32^. We serially transplanted total marrows from *drl:MYC:mCherry* donors into sub-lethally irradiated recipients and observed robust and rapid engraftment and disease propagation (7/7 recipients upon primary and 17/18 recipients upon secondary transplant, **Fig. S2**). In conclusion, overexpression of *MYC* led to a robust erythroleukemia in adult zebrafish and we used this model for further studies of clonality.

**Fig. 1.**
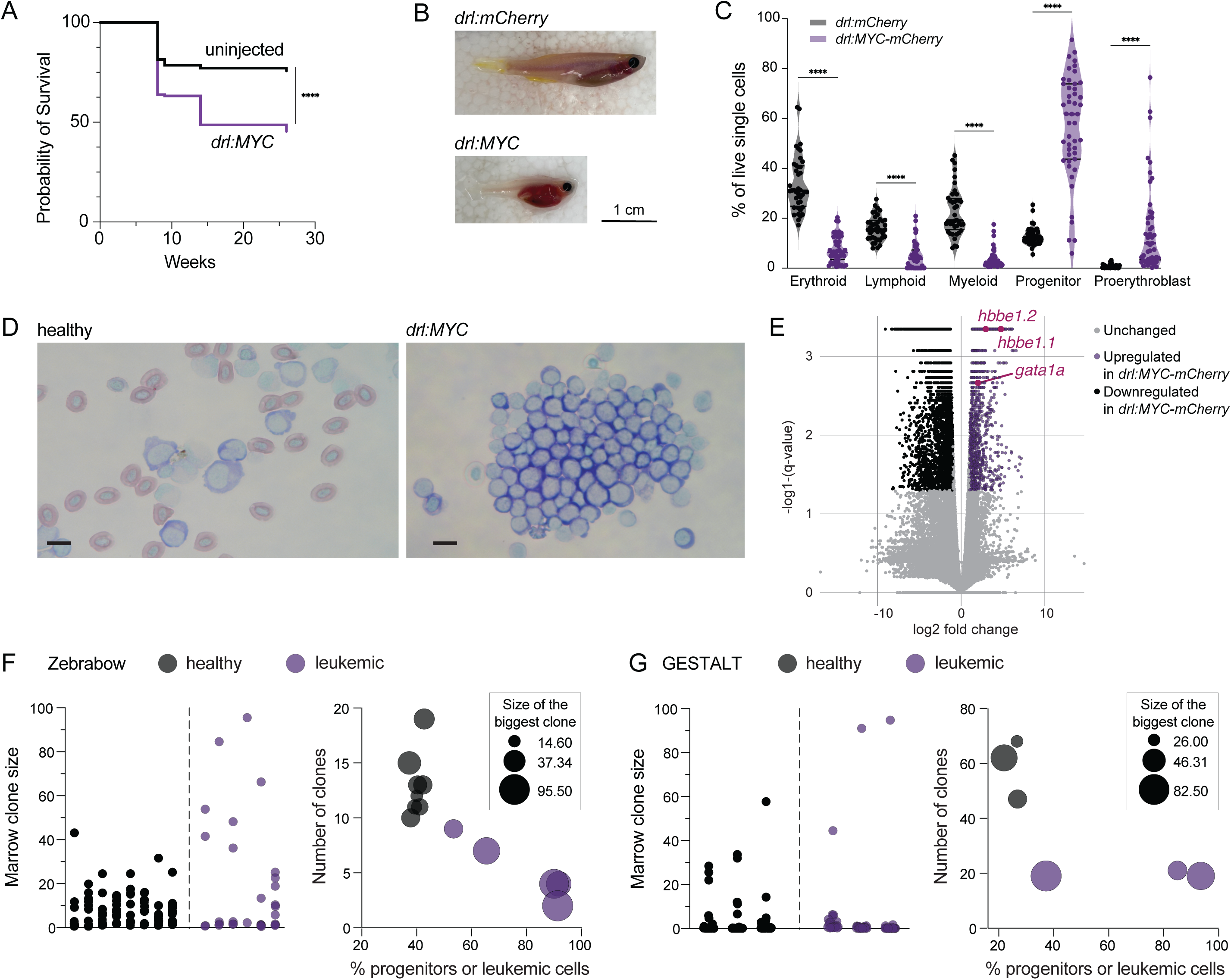
Overexpression of MYC induces acute myeloid leukemia. (A) Survival curves of uninjected (n=70) and *drl:MYC* injected (n=152) zebrafish demonstrates significantly reduced survival of *drl:MYC* animals. ****p<0.0001 (B) *drl:mCherry* control and *drl:MYC* leukemic animals in Casper EKK background demonstrated phenotypic manifestations of leukemic. Scale represents 1 centimeter. (C) Violin plot of live marrow cells quantified by flow cytometry and gated into five distinct populations as defined by forward and side scatter (FSC and SSC) demonstrating changes in hematopoietic lineages in *drl:MYC* marrows (n=40 *drl:mCherry*, n=48 *drl:MYC-mCherry,* ****p<0.0001). (D) May-Grünwald Giemsa staining of cytospins of total healthy and *drl:MYC* marrows demonstrating accumulation of leukemic cells in *drl:MYC* marrows. Scale bar 10μm (E) Violin plot with all genes differentially expressed between *drl:mCherry* control (n=2) and *drl:MYC* (n=4) total marrow cells. Purple dots are significantly upregulated in *drl:MYC* marrows and black dots are significantly downregulated in *drl:MYC* marrows with p-value cutoff < 0.05 and log2 fold change < −1 and > 1. (F) Zebrabow color barcoding of healthy (n=8) and leukemic (n=6) marrows. Left panel shows single progenitor or leukemic clone size in individual healthy and leukemic animals, respectively. Right panel shows the number of clones, % of healthy progenitors or leukemic cells as quantified by flow cytometry and size of the biggest clones demonstrating that higher leukemic burden is reflected by lower clone number and largest dominant clone. (G) GESTALT barcoding of healthy (n=3) and leukemic (n=3) marrows. Left panel shows single progenitor or leukemic clone size in individual healthy and leukemic animals, respectively. Right panel shows a visualization of number of clones, % of healthy progenitors or leukemic cells as quantified by flow cytometry and size of the biggest clones demonstrating that higher leukemic burden is reflected by lower clone number and largest dominant clone.

We used two independent clone tracing methods, Zebrabow and GESTALT, to probe healthy progenitor and leukemic cells. Zebrabow is a color-labeling based approach that allows for flow cytometric analysis of gated healthy progenitor and leukemic cell clones^33^. Most leukemic cells were composed of 1 or 2 dominant clone(s), contrasting with polyclonal healthy progenitors with 10 or more clones (**Fig. 1F**, left panel). Furthermore, leukemic marrows with fewer clones were associated with a higher disease burden, suggesting eventual outgrowth of the leukemic clone over non-mutant hematopoiesis (**Fig. 1F**, right panel). GESTALT is a CRISPR Cas9-based genetic lineage tracing method in which DNA barcodes are introduced in an artificial cassette during embryonic development and can be sequenced in adult sorted blood populations^27,28^. We sequenced GESTALT barcodes from sorted heathy progenitor and leukemic cells. We detected 33 to 68 total clones in healthy progenitors and 19 to 21 clones in leukemic cells with a single dominant clone in leukemic cells compared to healthy progenitors (**Fig. 1G**). The higher clone number detected with GESTALT compared to Zebrabow is expected due to the higher complexity of DNA barcodes compared to color barcodes. Overall, these orthogonal methods revealed that mosaic overexpression of human *MYC* in blood cells results in clonal expansion of 1 or 2 progenitors leading to leukemic transformation.

### Leukemia induces clonal selection of niche endothelial and stromal cells

We next aimed to probe the clonality of niche ECs and MSCs using the GESTALT system. We generated a GESTALT barcode transgenic line that carries the endothelial reporter *kdrl:GFP* and the stromal reporter *cxcl12a:dsRed* (**Fig. 2A**). We crossed this line to the GESTALT guide line and injected resulting 1-cell stage embryos expressing guide RNA (gRNA) targeting sites 1 to 4 of the GESTALT cassette and *cas9* mRNA to induce dynamic barcoding in the first four sites of the cassette during early embryogenesis. The *drl:MYC* construct or control construct was injected to induce leukemia in adults. Embryos were heat shocked at 28 hours post fertilization (hpf) to induce Cas9 expression and triggering barcodes in sites 5 through 9 of the cassette at the time of HSC birth. Barcoded adults were raised to adulthood and monitored for disease onset. Zebrafish with more than 60% of leukemic cells in the marrows were retained for analysis and matched to healthy control samples. Endothelial and stromal populations were sorted for DNA extraction, GESTALT cassette amplification and next-generation sequencing (**Fig. S3A**). We found that leukemic marrows had significantly decreased EC and MSC clone numbers compared to healthy controls indicative of a clonal selection (**Fig. 2B**). Clone size analysis revealed that leukemic endothelial and stromal clones were significantly larger than healthy clones suggesting a niche clonal expansion (**Fig. 2C**; **Fig. S3B,C**, left panels). We found a significant increase in median endothelial clone size suggesting an important endothelial clonal remodeling in leukemia marrows (**Fig. S3B,C**, right panels). The five largest clones had a significantly larger average size in leukemic marrows emphasizing a leukemia-driven niche selective clonal expansion (**Fig. 2D**). Overall, our data revealed that leukemia induced a significant clonal redistribution with clonal selection within the niche cells.

**Fig. 2.**
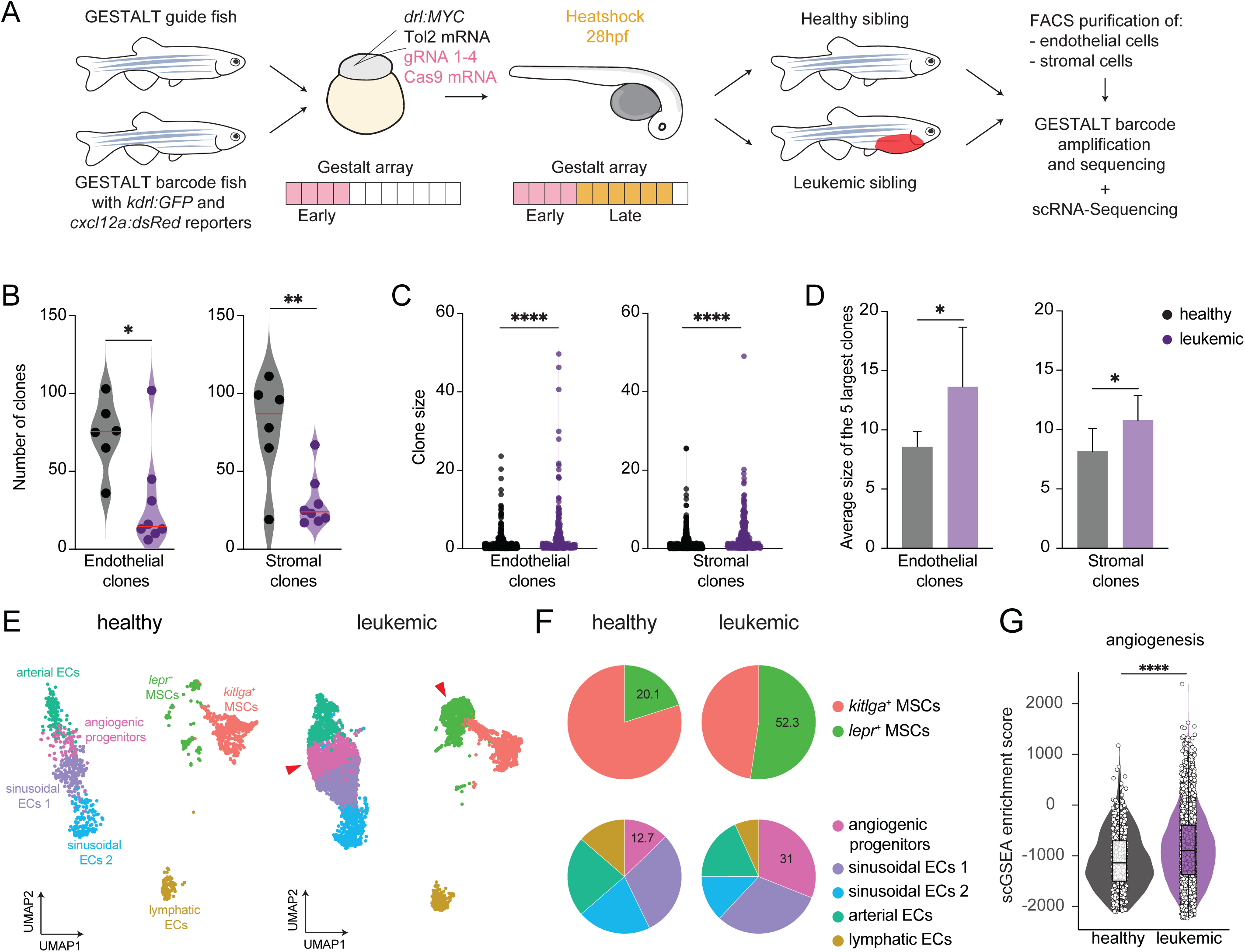
Leukemia induces clonal and transcriptional remodeling of niche ECs and ECs. (A) Experimental approach for GESTALT clone tracing of niche cells in leukemic zebrafish. GESTALT guide and barcode fish are crossed and embryos are injected with gRNA1-4 for early barcoding and *drl:MYC*. Embryos are heatshock at 28hpf for late barcoding. Adult zebrafish with leukemia are identified based on phenotypic manifestation of disease and siblings identified as *drl:MYC*^-^ by PCR on blood sample are included as controls. Marrows are dissected for sorting of *kdrl:GFP*^+^ ECs and *cxcl12a:dsRed*^+^ MSCs for DNA extractions GESTALT barcode amplification and sequencing and single-cell mRNA-sequencing. (B) Violin plot of the number of clones detected in purified ECs (left panel) and MSCs (right panel) in healthy (n=6) and leukemic marrows (n=8) showing decreased clonal complexity upon leukemia. * p<0.05, ** p<0.01. The red line indicates the median value. (C) Violin plot of EC (left panel) and MSC (right panel) clone size of pooled healthy and leukemic marrows revealing clonal selection upon leukemia. **** p<0.0001. (D) Bar graph of the mean with standard deviation of the size of the top five largest EC (left panel) and MSC (right panel) clones in healthy and leukemic marrows showing selective clonal expansion in leukemic marrows. * p<0.05. (E) UMAP dimensionality reduction of scRNA-seq of *kdrl:GFP*^+^ ECs and *cxcl12a:dsRed*^+^ MSCs from healthy (n=2) and leukemic (n=3) marrows (n=9,176 cells). Colors represent cell types and match the legend in panel F. (F) Pie chart of MSC (upper panel) and ECs (lower panel) sub-population proportions in healthy and leukemic marrows as identified in panel E. (G) Violin plot showing the angiogenesis scGSEA enrichment score in healthy and leukemic sinusoidal ECs demonstrated increase angiogenic activity of ECs in leukemic marrows. **** p<0.0001.

### Clonally selected *lepr*^+^ mesenchymal stromal cells have a unique gene expression signature in leukemic marrows

We reasoned that clonal remodeling of the niche is accompanied by transcriptional remodeling and functional change. To understand molecular changes in niche cells, we performed scRNA-seq of purified *kdrl:GFP*^+^ ECs and *cxcl12a:dsRed*^+^ MSCs from healthy and leukemic marrows. Combined data analysis identified 7 sub-populations (**Fig. 2E**). Two MSC subpopulations could be identified based on the differential expression of the niche markers *kitlga* and *lepr* (**Fig. S4A**, **Supplementary Table 1**). Both *kitlga^+^* and *lepr^+^* MSC proportions were altered upon leukemia with an expansion of the *lepr^+^* MSC subpopulation (**Fig. 2F**; **Fig. S5A**). To further understand the transcriptional changes in these populations, we performed single-cell GSEA analysis between healthy and leukemic MSCs. This analysis revealed a significant increase of IL6/JAK/STAT3, TGF beta signaling and interferon alpha response (**Fig. S5B**). Differential gene expression analysis identified the stromal cell activation marker *cd248a* expressed exclusively in *lepr^+^* MSCs and upregulated in leukemic marrow (**Fig. S5C**). Immunohistochemistry for dsRed in healthy marrows revealed an even distribution of *cxcl12a:dsRed*^+^ MSCs (**Fig. S5D**). In leukemic marrows, two distinct MSC regions could be observed: MSC-depleted regions with rare MSCs packed between leukemic cells and MSC-dense regions with a higher density of MSCs. Our clonal, transcriptional and imaging analysis of MSCs in leukemic marrows suggested a spatially restricted clonal activation of *lepr^+^* MSC.

### Endothelial progenitors are expanded and drive spatially defined angiogenesis in leukemic marrows

Our prior work demonstrated that sinusoidal ECs intimately interact with HSCs and drive normal stem cell retention and division in the niche^14^. We have also demonstrated that endothelial fates drive ectopic hematopoietic niche formation^34^. To define the endothelial populations present in the zebrafish leukemic marrow, we used our scRNA-seq data to identify arterial, lymphatics and sinusoidal ECs, with sinusoidal ECs further subdivided into three distinct subpopulations (**Fig. S4B**, **Supplementary Table 1**). This included two subpopulations of sinusoidal ECs, with subpopulation 2 expressing marker genes reported as niche factors in the zebrafish embryonic stem cell niche^34^ and subpopulation 1 expressing several angiogenic factors and sharing marker genes with arterial and sinusoidal ECs (**Fig. S6A**). Trajectory analysis using Monocle3 supported the notion that these angiogenic progenitors differentiate towards arterial and sinusoidal ECs (**Fig. S6B**)^35^. EC subpopulations were altered in leukemic marrows with a higher proportion of angiogenic progenitors (**Fig. 2F**). Live imaging of *kdrl:GFP* healthy and leukemic marrows showed a change in spatial organization of the vasculature in leukemia. In healthy marrows, *kdrl:GFP* cells were evenly distributed forming complex sinusoidal networks (**Fig. S6C**). In leukemic marrows, regions of dense sinusoidal vasculature could be identified (**Fig. S6C**, red arrows) with EC exhibiting high endosomal content, a cellular requirement for angiogenesis^36^(**Fig. S6D**). This finding was validated by higher expression of endosome markers in leukemic marrows with scRNA-seq (**Fig. S6E**). Single-cell GSEA analysis between healthy and leukemic sinusoidal ECs revealed a significant enrichment of angiogenesis-related genes (**Fig. 2G**). Altogether, GESTALT, scRNA-seq and imaging analyses suggest that leukemia induces clonal angiogenesis via activation of angiogenic progenitors to generate spatially altered sinusoidal networks.

### The leukemia-secreted angiogenic factor *apelin* mediates endothelial cell clonal expansion

We hypothesized that leukemic cells must produce a factor to induce angiogenic progenitor activation and sinusoidal EC clonal expansion to promote disease progression. We performed scRNA-seq of healthy hematopoietic and leukemic cells to identify ligand(s) significantly upregulated by leukemic cells that had paired receptor(s) upregulated on leukemic ECs. We identified three candidates: vascular endothelial growth factor c (*vegfc*), adrenomedullin b (*admb*) and apelin (*apln*) (**Fig. 3A,B**; **Fig. S7A,B**). We generated *draculin*-driven overexpression constructs for these three candidates and injected them into GESTALT embryos to test their individual ability to drive niche clonal and transcriptional changes in the absence of leukemia. Overexpression of *vegfc* induced a moderate EC clonal expansion that stemmed from small clones without MSC clonal expansion, and overexpression of *admb* did not induce any clone expansion (**Fig. S7C,D**). The *apln* receptors are only expressed by ECs and *apln* overexpression resulted in a significant EC clonal expansion but no MSC clonal changes (**Fig. 3C**; **Fig. S7E,F**). To assess the transcriptional changes in niche cells upon *apln* overexpression, we performed scRNA-seq of purified *kdrl:GFP*^+^ ECs and *cxcl12a:dsRed*^+^ MSCs from healthy and *apln* overexpression marrows (**Fig. 3D**; **Fig. S4**). We found that *apln* overexpression induced an increase in angiogenic progenitors and differentiation into niche sinusoidal ECs similar to *drl:MYC* leukemia (**Fig. 3E**).

**Fig. 3.**
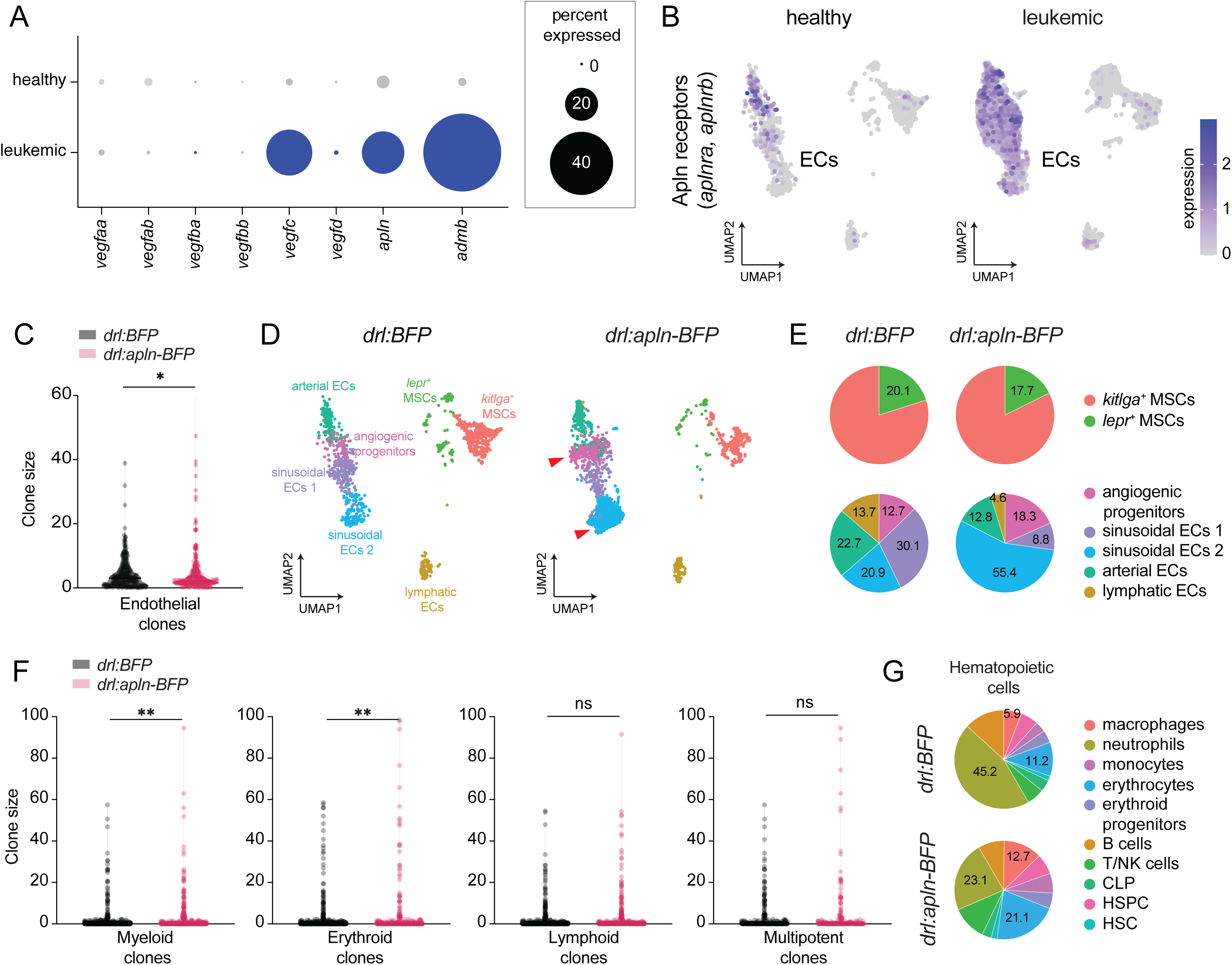
*apelin* secreted by blood cells regulates EC clonal and transcriptional profile. (A) Dot plot highlighting candidate secreted factors identified by DEG analysis between healthy HSPCs and leukemic cells. Dot size represents the fraction of cells expressing each gene. *vegfc*, *admb* and *apln* were selected for further functional analyses. (B) UMAP dimensionality reduction as defined in Fig. 2E. showing the combined expression of the two isoforms of the apln receptor: *aplnra* and *aplnrb* in healthy and leukemic marrows. Color represents gene expression level. The receptors are exclusively detected in ECs. (C) Violin plot of EC clone size of pooled *drl:BFP* control (n=12) and *drl:apln-BFP* (n=14) overexpressing marrows. * p<0.05. (D) UMAP dimensionality reduction of scRNA-seq of *kdrl:GFP*^+^ ECs and *cxcl12a:dsRed*^+^ MSCs from control (n=2) and *apln* overexpression (n=2) marrows. Colors represent cell types and match the legend in panel E. (E) Pie chart of MSC (upper panel) and ECs (lower panel) sub-population proportions in control and *apln* overexpressing marrows as identified in panel D. (F) Violin plot of myeloid, erythroid, multipotent and lymphoid clone size of pooled control (n=12) and *apln* overexpressing (n=14) marrows revealing clonal expansion of myeloid and erythroid clones upon overexpression of *apln* alone. ** p<0.01. (G) Pie chart of hematopoietic sub-population proportions in control and *apln* overexpressing marrows with increased macrophages and erythrocytes.

We then posited that overexpression of *apln* leads to increased production of ECs altering hematopoiesis. To test this, we examined the clonality and cell type output in control and *drl:apln-BFP* marrows. We found that overexpression of *apelin* significantly increased myeloid and erythroid clone size (**Fig. 3F**). Flow cytometric analysis of marrows showed no significant changes in the overall hematopoietic output (**Fig. S7G**). scRNA-seq of hematopoietic cells from control and *apln* overexpressing marrows revealed an increase in mature erythroid cells upon *apln* overexpression while the fraction of erythroid progenitors remains unchanged (**Fig. 3G**). Using transgenic reporter lines, we confirmed that the number of *gata1a:dsRed*^+^ erythroid progenitors was unchanged but significantly more *lcr:GFP*^+^ mature erythroid cells were present upon *apln* overexpression (**Fig. S7H**). scRNA-seq demonstrated an increase in the fraction of macrophages at the expense of neutrophils. We did not find a difference in the fraction and transcriptional state of HSCs and HSPCs and confirmed this result using the *runx1+23:mCherry* transgenic line which revealed no difference in HSPC number upon *apln* overexpression (**Fig. S7I**). Although HSCs did not show a clonal change or change in gene expression upon *apln* overexpression, the more sensitive clonal output of myeloid and erythroid cells (which in zebrafish maintain their nucleus) suggests HSC clonal output and differentiation changes to a myelo-erythroid fate via niche ECs.

### *apln-*mediated EC remodeling is necessary and sufficient to drive leukemogenesis

In our *apln* overexpression model, 7% of animals (5 out of 71) exhibited evidence of leukemic disease with a significant increase of progenitor-like cells with concomitant block in myeloid differentiation (**Fig. 4A**; **Fig. S8A**). May-Grünwald Giemsa staining of marrow cell cytospins revealed an abundance of dark purple blasts with high nuclear-to-cytoplasmic ratio, recapitulating the phenotype observed in *drl:MYC* leukemic animals (**Fig. 1D**, **4B**). Next, we asked if *apln* was required for MYC-driven leukemogenesis. To test this, we generated a stable *apln* loss-of-function zebrafish line. We crossed *apln* heterozygous animals and injected embryos with *drl:MYC*. Animals with confirmed *drl:MYC* transgene were followed for leukemia incidence. Compared to WT zebrafish, *apln*^het^ and *apln*^hom^ mutants had a dose-dependent lower incidence of leukemia (**Fig. 4C**). In fact, we did not observe any death due to leukemia in the *apln*^hom^ mutants at 25 weeks. These data showed that *apln* secreted by leukemia cells is necessary and sufficient to induce disease progression.

**Fig. 4.**
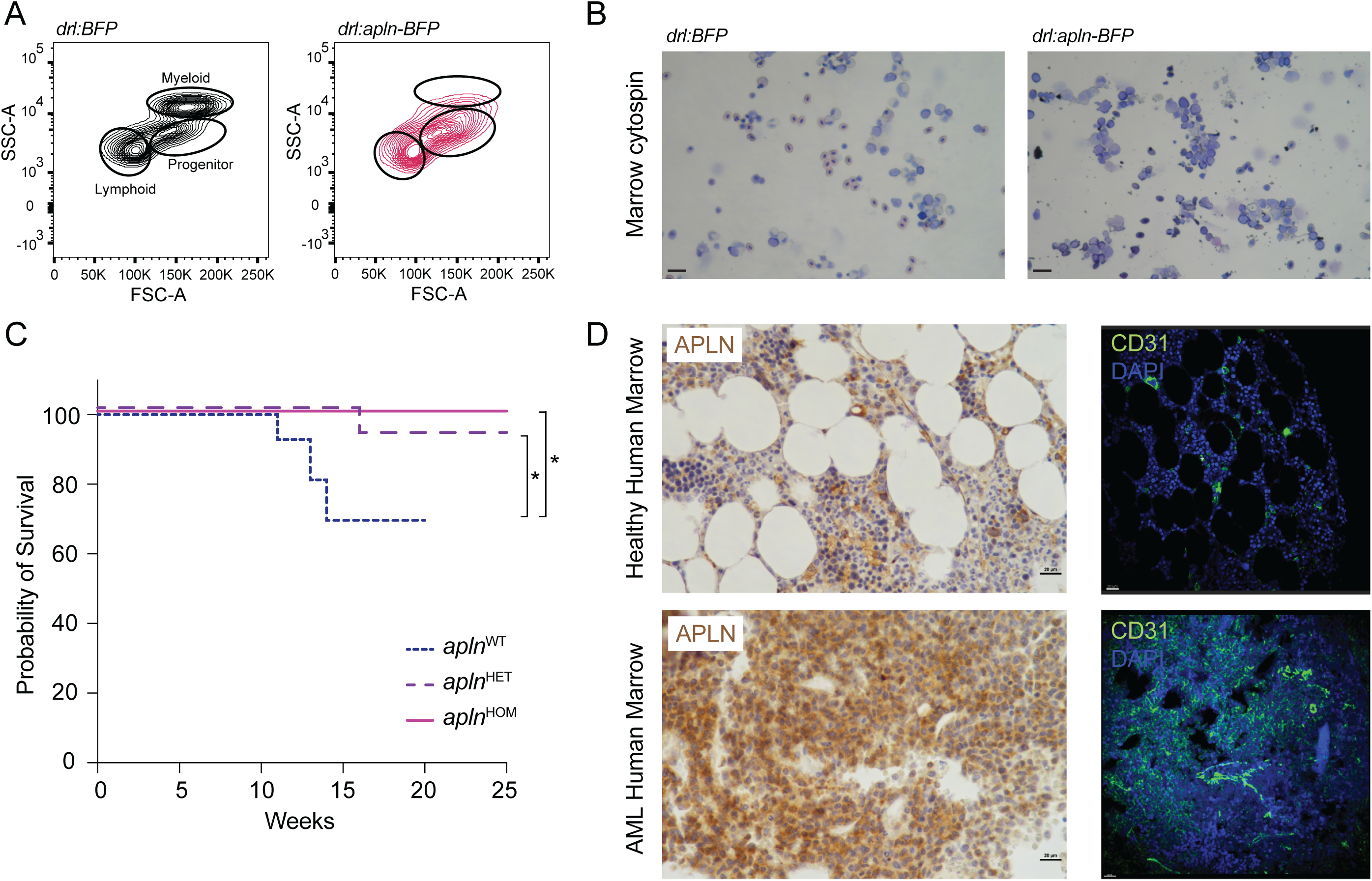
*apln* mediated EC remodeling drives leukemogenesis. (A) Flow cytometry contour plot of a representative example of *drl:BFP* control and *drl:apln-BFP* abnormal marrow showing a leukemia-like progenitor profile. (B) May-Grünwald Giemsa staining of cytospins of total control and *apln* overexpressing marrows demonstrating accumulation of leukemic cells in *drl:apln-BFP* marrows. Scale bar 20μm (all panels). (C) Survival curves of *apelin* wildtype (*apln*^wt^, n=6), heterozygous (*apln*^het^, n=9) and homozygous (*apln*^hom^, n=4) mutant animals injected with and positive for *drl:MYC* overexpression transgene. * p<0.05. (D) APLN immunohistochemistry (left panels) and CD31 immunohistochemistry (right panels) of one representative example of healthy human marrow and one representative example of AML human marrow. Scale bar 20μm (all panels).

Lastly, we asked if human AMLs express APLN. Our analysis of the BEAT AML dataset revealed a significant upregulation of APLN in leukemic samples compared to healthy bone marrow mononuclear cells (**Fig. S8B**)^37^. To validate this result, we performed immunohistochemistry for APLN and immunofluorescence for the EC marker CD31 on adjacent sections of healthy and AML marrows. All AML marrows had higher APLN staining compared to controls, although heterogeneity in expression was noted (**Fig. 4D**; **Fig. S8C,D**). CD31 staining revealed higher vessel density in APLN high-expressing AML marrows compared to control suggesting APLN-driven angiogenesis in human disease. These results indicated that human AML cells produced high levels of APLN and that the marrow vasculature is affected in these patients.

## Discussion

We evaluated the clonality of the leukemic niche *in vivo* using a novel zebrafish model of AML. We demonstrated that MYC-driven AML induces a clonal selection of niche ECs and MSCs. We identified the leukemia-secreted angiogenic peptide *apln* as responsible for EC clonal expansion and reprogramming towards an EC progenitor state driving angiogenesis. By genetically overexpressing *apln* in absence of MYC, we demonstrated niche EC clonal selection is associated with an alteration of HSCs clonal output in the myelo-erythroid lineage. *Apelin* independently was able to drive leukemogenesis, albeit with reduced penetrance. Genetic ablation of *apln* reduced the incidence of leukemia in a short follow-up period in our *drl:MYC* leukemia model. This supports the proposed cellular mechanism of leukemic cells remodeling the endothelial niche via the secretion of a single ligand which will reciprocally promotes leukemic cell growth. The role of the marrow niche in hematological disorders has been demonstrated by introducing mutations solely in niche compartments which resulted in MDS and/or pre-leukemic states^4–8^. Our work demonstrates for the first time that a secreted ligand inducing niche remodeling can promote malignant transformation, emphasizing the role of the niche in disease onset and progression.

The VEGF family is the most characterized family of secreted pro-angiogenic factors that have been shown to promote vascular growth and remodeling and its levels have been shown to be increased in AML patients^38,39^. We found that ap*ln* is more potent to trigger EC clonal selection than *vegfc*, the only VEGF family member we found secreted by leukemic cells *in vivo* in our leukemia model. Clinical trials testing anti-VEGF therapy were terminated due to lack of efficiency and our work places *APLN* signaling as a prime candidate for clinical intervention^40,41^. We demonstrated the involvement of APLN signaling in human disease by probing for the expression of APLN in AML marrows by immunohistochemistry and observing a strong increase in APLN levels in tandem with increase vessel density as measured by CD31 expression. The relevance of APLN in human leukemias is supported by the identification of a risk variant in the APLNR gene in a Genome-wide association study (GWAS) of pediatric acute lymphoid leukemia patients^42^. While anti-VEGF therapy may still have an important role but developing antibody therapy to APLN could help treat leukemia. Other growth factors may regulate niche clonality in different types of leukemia and tumors. Additionally, growth factors that drive MSC clonal changes remain to be uncovered. We anticipate that our work will initiate studies centered on targeting niche clonality in cancer.

## Acknowledgments

We thank the aquatics research staff at Boston Children’s Hospital (BCH), the BCH flow cytometry core and the Harvard Bauer Core for next-generation Sequencing. We thank Dana-Farber/Harvard Cancer Center in Boston, MA, especially Caitlin Edwards, for the use of the Specialized Histopathology Core, which provided histology and immunohistochemistry service. Dana-Farber/Harvard Cancer Center is supported in part by an NCI Cancer Center Support Grant # NIH 5 P30 CA06516. We thank all past and present Zon lab members for critical feedback on the research and the manuscript, especially Dr. Samuel Wattrus, Dr. Brandon Gheller, Dr Kyle Drake and Dr Rohan Bhattacharya.

## Funding

National Institutes of Health grant 1 R01 DK140372-01 (LIZ)

National Institutes of Health grant RC2 DK120535 (LIZ)

National Institutes of Health grant 2 P01 HL131477-06A1 (LIZ)

Edward P. Evans Foundation (LIZ)

Alex’s Lemonade Stand Fund, Crazy 8 Initiative Award Program (LIZ, SA, JS)

European Molecular Biology Organization Postdoctoral Fellowship LT_290_2019 (CSB)

Human Frontiers Science Program Postdoctoral Fellowship LT000494/2020-L (CSB)

National Institutes of Health grant 1 K99 HDK134760 (CSB)

National Institutes of Health grant P20GM103423 (R. Menard, R. Madelaine)

National Institutes of Health grant P20GM104318 (R. Menard, R. Madelaine)

National Institutes of Health grant P20GM144265 (R. Menard, R. Madelaine)

National Institutes of Health grant DP2GM149750-01 (AM)

Pew Charitable Trust (AM)

LIZ and JS are Howard Hughes Medical Institute Investigators.

## Author contributions

CSB and LIZ conceptualized the study. CSB, OM, SA, SY, ALR, RP, R. Menard and AM performed experiments and analyses. CSB wrote the original draft which was reviewed and edited by SA, R. Madelaine, AM and LIZ.

## Data and materials availability

The bulk RNA-seq, scRNA-seq and GESTAL barcode data is currently being deposited in GEO and accession number with be provided shortly. Code used for data analysis in available upon request.

## Materials & Correspondence

Dr. Leonard Zon.

## Materials and methods

### Animal models

Wild-type zebrafish *Tübingen* (*TU*), *Casper EKK*, and transgenic lines GESTALT barcode and guide lines^28^, *draculin:CreER^T2^*^29^, *Zebrabow-M*^43^, *kdrl:GFP*^44^, *cxcl12a:dsRed2*^45^, *Runx1+23:mCherry*^14^, *gata1a:dsRed*^30^ and *lcr:GFP*^31^ were used in this study. All animals were housed at Boston Children’s Hospital and handled according to approved Institutional Animal Care and Use Committee (IACUC) of Boston Children’s Hospital protocols.

### Transgenesis

Overexpression constructs were generated by DNA synthesis of desired cDNA (MYC, *apln*, *vegfc*, *admb*), D-TOPO cloning into pME Gateway, and LR Gateway Reaction to generate constructs with the *draculin* promoter driving a T2A fused with *TagBFP*, *mCherry* or *GFP*, followed by a SV40 polyA signal, all flanked by Tol2 integration sites. The fidelity of all constructs was confirmed by sequencing prior to injection. Constructs were injected in embryos at a concentration ranging from 5 to 25 ng/ul (determined experimentally to maximize embryo survival and mosaicism) with Tol2 mRNA at 25 ng/ul in the injection mix.

### Zebrafish apelin mutant generation

To generate the *apelin* mutant, we took advantage of a previously described genome-editing method^46^. Specifically, we used tracrRNA and crRNA (IDT) to form functional gRNA duplexes targeting the ORF of the apelin gene (5’-GAATGTGAAGATCTTGACGC-3’). Co-injection of the gRNAs with the Cas-9 protein (IDT) resulted in indels in the apln gene coding sequence, identified by Sanger sequencing and analyzed with ICE software (Synthego) in F0 injected embryos. F0 carriers for mutations in the apelin locus in the germline were identified by sequencing the alleles on the clutches and were then outcrossed to obtain F1 heterozygous mutants. Mutations (indels) leading to a premature stop codon were identified by ICE analysis (Synthego) of the Sanger sequencing results. F2 *apln* homozygous mutants were also identified by sequencing. The apelin mutant harbors a 26-bp insertion leading to a stop codon at position 18 of the peptide.

### Leukemic cell transplants

Recipients were 3 months old Casper EKK zebrafish irradiated on days −2 and −1 with 14-Gy each day for a total of 28-Gy sublethal dose. Donors were two *drl:MYC-mCherry* animals exhibiting a leukemic phenotype. On day 0, each donor was dissected and marrows were collected as described below. Six WT TU marrows were collected to serve as helper marrows. A small fraction of cells from leukemic single cell suspension from both donors were taken for flow cytometry analysis to confirm the presence of *mCherry*+ leukemic cells prior to transplantation. The rest of the marrow suspensions were counted then washed twice with 1X PBS. Helper marrow cells were pooled and divided over all recipients. 2.5 μl of cells (mix of leukemic and helper cells) were transplanted retro-orbitally in each recipient. For donor 1, 3,165,000 cells were transplanted per recipient (n=4) and for donor 2 4,650,000 cells were transplanted per recipient (n=3). Recipients were monitored weekly for phenotypical manifestation of leukemia and were sacrificed for analysis prior to disease-induced death (within 5 to 10 weeks post-transplant). Marrows from recipients were dissected and analyzed by flow cytometry and engraftment was defined as ≥2% mCherry+ cells or ≥25% cells in the progenitor gate (as defined by FSC/SSC in WT TU marrows). Cells from engrafted recipients from both primary donors were counted and transplanted in irradiated secondary Casper EKK recipients following the same procedure as described above. For donor 1, 591,000, 594,000, 555,000 and 198,000 cells were transplanted in secondary recipients (n=6). For donor 2, 323,000, 1,395,000 and 602,000 cells were transplanted in secondary recipients (n=13). All secondary recipients exhibited signs of disease within 6 weeks and were analyzed as described above.

### Zebrabow color labeling

Zebrabow-M adults were crossed with *draculin:CreER^T2^*adult and embryos were injected with *drl:MYC* and *Tol2* mRNA. At 28 hours post fertilization (hpf) embryos were transferred to 6-well plates at a density of 25-35 embryos per well and treated with 15 μM 4-hydroxytamoxifen (4-OHT) for 3-5 hours in the dark at 28.5°C.

### GESTALT embryo barcoding

Barcoding was performed as described previsouly in ^28^. Briefly, GESTALT barcode females were crossed with a GESTALT guide male to minimize maternal inheritance of gRNA 5-9 (**Fig. 2A**). Single cell embryos were injected with gRNA 1-4 (each gRNA at 17.5 ng/μl in injection mix) and cas9 mRNA (150 ng/μl in injection mix). Overexpression constructs were included in the injection mix as appropriate. At 28 hpf, embryos were transferred to 1.5 ml tubes at a density of 25-35 embryos in 500 μl of embryo water and heated at 37°C for 30 minutes.

### GESTALT barcoded zebrafish genotyping

At 2 months post fertilization, all barcoded animals were fin clipped and DNA from fin samples was extracted (Zymo Quick DNA miniprep). The GESTALT cassette was amplified by PCR using Phusion High-Fidelity PCR Master Mix with HF Buffer (NEB) with the following conditions: 98°C 3 minutes, [98°C 10 seconds, T annealing = 63°C 10 seconds, 72°C 10 seconds] x 35 cycles, 72°C 5 minutes and using primers FW: CTGCCATTTGTCTCGAGGTC and RV: CTGCCATTTGTCTCGAGGTC. The editing level was assess using 1) gel electrophoresis to detect the presence of a smear below the 310 base pairs (bp) unedited cassette band and 2) next generation sequencing for samples with a smear detected on gel. Reads were mapped using the pipeline described below. Animals with 75% and above edited reads were selected for further analysis.

### Marrow dissection and flow cytometry

Zebrafish were euthanized according to IACUC guidelines and kidney marrows were dissected under a Leica MZ75 light microscope. For hematopoietic cell collection, tissue was collected in cold 1X DPBS (Gibco) with 2% fetal bovine serum (FBS, Gemini Bio-Products) and 1 USP units/mL heparin (Sigma) (blood buffer), and then mechanically dissociated by repeated pipetting and filtered through a 40-μm nylon mesh prior to adding 3 nM of DRAQ-7 (Abcam) for viability assessment for flow cytometry analysis and/or sorting. For EC and MSC recovery, tissue was collected in cold 1X PBS and incubated in Liberase (Roche) for 25 minutes at 37°C with 600rpm agitation and mechanical dissociation using a p200 pipet tip after 10 and 20 minutes. Pure FCS was added to reach 10% volume and stop the enzymatic reaction. Cell suspension was filtered through a 40-μm nylon mesh and washed with 1ml of cold 1X DPBS with 2% FBS at 400g for 5 minutes. Pellet was resuspended in cold 1X DPBS with 2% FBS with 3nM of DRAQ-7 for flow cytometry analysis and/or sorting. Flow cytometric analysis and sorting was performed on a BD FACSAria II or FACSFortessa (BD Biosciences). Gates were drawn using negative controls and all data was analyzed using FlowJo. For all animals, two types of unsorted samples were kept for DNA extraction 1) an aliquot of 10,000-50,000 total unsorted marrow cells 2) a peripheral blood sample collected by cardiac aspiration using a p10 tip coated with heparin and deposited in 300 μl of cold blood buffer and filtered through a 40-μm nylon mesh.

### Adult Zebrabow color analysis

Color barcodes from Zebrabow kidney marrow samples were quantified using previously published pipelines^33^ adapted to a Python-based interface. The progenitor (as defined by FSC/SSC gating) color output was chosen as a read out of clonal changes.

### Single cell RNA-Sequencing library preparation

Single cells gated for targeted cell populations (lymphoid, myeloid, progenitor gates as defined by FSC/SSC or *kdrl:GFP*^+^ and *cxcl12a:dsRed*^+^ ECs and MSCs, respectively) were sorted in 300 μl of PBS supplemented with 0.5% Bovine Serum Albumin (Gemini Bio-Products). Sorted cell numbers were ranging between 18,000 and 100,000 cells. Samples were centrifugated at 500g for 5 minutes and cells were resuspended in 30 μl for cell counting using a hemocytometer. Live cell concentrations were assessed and we aimed for a targeted recovery of 5,000 to 8,000 cells per lane using the 10X Genomics Chromium Next GEM Single Cell 3’ Reagent Kit v3.1. Sequencing libraries were generated following manufacturer’s instructions and amplified cDNA and final libraries quality was probed using high sensitivity DNA D5000 and D1000 tape station kits (Agilent). Libraries were sequenced on NovaSeq platform (SP flow cell) with a targeted minimum of 20,000 paired end reads per cell.

### Single cell RNA-Sequencing data analysis

10x genomic scRNA-seq data was analyzed by 10x CellRanger v7.0 package. The sequencing reads were aligned to the zebrafish genome Ensembl GRCz11. The CellRanger count command with –include-introns option generates gene-barcode matrix for each sample, which were imported into R using the Seurat suite version 3.0^47–50^. Briefly, low quality cells were filtered out by selecting cells with the following parameters: nFeature_RNA > 600 & nFeature_RNA < 5000 & percent.mt < 12. The filtered samples were integrated across samples groups using the 5000 most variable genes as anchors. PCA dimensionality reduction using 30 PCs was performed, followed by clustering and Uniform Manifold Approximation and Projection (UMAP) using a value of 0.4 for the resolution. Downstream differential gene expression (DGE) analyses were performed with the FindMarkers() functions. Cell type signatures were generated using the AddModuleScore() function based on unbiased DGE analyses and curated lists from published literature^51^. For single-cell GSEA analysis, the Escape package was used ^52^ with the Hallmark gene set from the Human Molecular Signatures Database (MSigDB). For Monocle 3, the SeuratWrappers package was used to create a Monocle object from a Seurat object^35,53,54^. Downstream analysis of ECs was performed using the cluster of angiogenic progenitors as root. Pseudotime values were added to the Seurat object and plotted using the DimPlot() function.

### GESTALT barcode library preparation

Single cells gated for targeted cell populations (lymphoid, myeloid, progenitor gates as defined by FSC/SSC or *kdrl:GFP*^+^ and *cxcl12a:dsRed*^+^ ECs and MSCs, respectively) were sorted in 300 μl of cold 1X DPBS (Gibco) with 2% fetal bovine serum (FBS, Gemini Bio-Products) and 1 USP units/mL heparin (Sigma). At least 2,000 single cells were sorted for each individual sample and animals for which at least one cell population did not reach 2,000 viable sorted cells were excluded for further analysis. Sorted cells were spun down and DNA was extracted (Zymo Quick DNA miniprep). GESTALT cassette amplification was performed using Phusion High-Fidelity PCR Master Mix with HF Buffer (NEB) with the following conditions: 98°C 3 minutes, [98°C 10 seconds, T annealing = 63°C 10 seconds, 72°C 10 seconds] x 35 cycles, 72°C 5 minutes and using primers FW: CTGCCATTTGTCTCGAGGTC and RV: CTGCCATTTGTCTCGAGGTC. PCR amplicons were purified on Qiagen MinElute or PCR purification kit columns prior to sequencing. An aliquot of the purified sample was run on gel to confirm purity and the presence of a smear below 310bp to confirm editing. Sequencing was performed on the Illumina MiSeq.

### GESTALT barcode data analysis

DNA sequencing amplicons were aligned and collapsed by unique molecular identifier using a modified version of our previous pipelines^27^. Insertion and deletion events were called at each target site, generating a set of target site calls for each unique molecule, and statistics files on each collapsed read were generated. The modified pipeline and associated code are available on GitHub here: https://github.com/aaronmck/Zon_zebrafish. Mapped read count files were loaded into R Studio for downstream analysis. Samples were blinded regarding their healthy and leukemic status for the analysis that follows. Briefly, barcodes from all samples in this manuscript were collided and frequency of barcodes were plotted. Barcodes detected in more than 25% of samples were considered common barcodes and excluded for further analysis. For each zebrafish, barcodes from all samples were merged and only barcodes with more than 10 mapped reads were considered for clonality analyses. For niche analyses, blood barcodes were identified from unsorted peripheral blood samples and subtracted from endothelial and stromal samples to exclude potential red blood cell contamination. Overlap between endothelial/stromal barcodes and other sorted hematopoietic population (lymphoid, myeloid and progenitors based on FSC/SSC gating) was examined and hematopoietic barcodes were removed from endothelial and stromal barcodes when detected at a frequency lower than 10%. This approach excluded all hematopoietic clones from stromal samples. However, this approach allows for preservation of a small fraction of barcodes in endothelial samples, likely reflective of the endothelial origin of the hematopoietic system. Only barcodes unique to endothelial, stromal or hematopoietic populations were kept for further quantification of barcode/clone numbers and clone fraction were calculated for each sample. For hematopoietic lineage analysis, multipotent clones are defined as clones detected in erythroid (peripheral blood), lymphoid and myeloid gates. Erythroid clones are defined as clones detected in the peripheral blood sample. Lymphoid and myeloid clones are defined as clones detected in their sorted respective gates based on FSC/SSC gating. Erythroid, myeloid and lymphoid clones include clones unique to each lineage and multipotent clones.

### Bulk RNA-Seq library generation

Total marrow cells were pelleted at 400g for 5 minutes and resuspended and vortexed in RLT buffer (Qiagen) supplemented with 1% beta-mercaptoethanol prior to storage at −80°C. RNA was extracted using a RNeasy Plus Micro Kit (Qiagen). 100 ng of RNA was processed using the RiboGone-Mammalian kit (Takara) followed by amplified cDNA generation using the SMARTer Universal Low Input RNA kit (Takara). cDNA quality was assessed using high sensitivity DNA D1000 tape station kits (Agilent). Sequencing libraries were generated using the ThruPLEX DNA-Seq Kit (Takara) and quality was assessed using high sensitivity DNA D1000 tape station kits (Agilent).

### Bulk RNA-Seq sequencing and data analysis

Libraries were sequenced on an Illumina Hiseq-4000. Quality control of RNA-Seq datasets was performed by FastQC and Cutadapt to remove adaptor sequences and low-quality regions. The high-quality reads were aligned to Ensembl GRCz11 of zebrafish genome using STAR 2.7.0 Spliced Transcripts Alignment tool^55^. The raw read counts of each gene is calculated by HTSeq^56^. Downstream analyses were performed using R Studio.

### Cytospin

100-200 μl of total single cell suspensions from marrow samples were loaded on cytology funnels with filter cards (Fisherbrand) and spun down at 500rpm for 5 minutes at medium acceleration in Shandon Cytospin Centrifuge onto glass slides. Slides were immediately stained using May-Grünwalk Giemsa stain for 4 minutes followed by two washes in deionized water (4 minutes each). Slides were air dried at room temperature over-night prior to imaging using a Nikon Eclipse E600 equipped with a DS-Ri3 camera. All images were acquired and processed using NIS-Elements (Nikon).

### Immunohistochemistry

Zebrafish were euthanized according to IACUC guidelines and fixed in 10% neutral Buffered Formalin (VWR) for 24 hours at room temperature. Immunohistochemistry was performed on the Leica Bond III automated staining platform using the Leica Biosystems Refine Detection Kit (Leica; DS9800). FFPE tissue sections were baked for 30 minutes at 60°C and deparaffinized (Leica AR9222) prior to staining. Primary antibodies were incubated for 30 minutes, visualized via DAB, and counterstained with hematoxylin (Leica DS9800). The slides were rehydrated in graded alcohol and cover slipped using the HistoreCore Spectra CV mounting medium (Leica 3801733).

The following antibodies were used:

- Anti mCherry (Abcam ab167453, 1:600 with a 20M EDTA antigen retrieval (Leica ER2 AR9640)).
- Anti GFP (Cell Signaling Technology 2956, clone D5.1, 1:50 with a 20M EDTA antigen retrieval (Leica ER2 AR9640)).
- Anti APELIN (Abcam, ab125213, 1:100 with a 20M EDTA antigen retrieval (Leica ER2 AR9640)).

### Immunofluorescence

Immunofluorescent staining was performed on the Leica Bond RX automated staining platform using the Leica Biosystems Refine Detection Kit (Leica DS9800). FFPE tissue sections were baked for 30 minutes at 60°C and deparaffinized (Leica AR9222) prior to staining. Following staining, slides were counterstained with DAPI (Nucblue; Invitrogen R37606) and cover slipped (Prolong Diamond; Invitrogen P36961). The following antibody was used: anti CD31 (CST, 3528, clone 89C2, 1:1600, antigen retrieval Leica ER1 AR9961 for 30min).

### Live imaging of marrows

Zebrafish were euthanized according to IACUC guidelines and immediately dissected for marrow collection. Marrows were slowly collected in one piece and placed in a glass bottom 6-well plate. A drop of cold PBS was deposited above the tissue and a round coverslip was superposed to maintain the tissue in contact with the glass bottom. Live imaging microscopy was performed using a Yokogawa CSU-X1 spinning disk mounted on an inverted Nikon Eclipse Ti microscope equipped with dual Andor iXon EMCCD cameras and a motorized x-y stage to facilitate tiling and imaging of multiple specimens simultaneously. All images were acquired using NIS-Elements (Nikon) and processed using Imaris (Bitplane).

### Statistics

Graphs and statistical analysis are done with R Studio (Posit) and Prism (GraphPad Software, Inc.). Quantitative graphs provide the mean and standard deviation or the median, as indicated in the figure legends. Statistical tests used were 2way ANOVA or t-test and p-values are indicated in the figure legends.

---

**Fig. S1.**
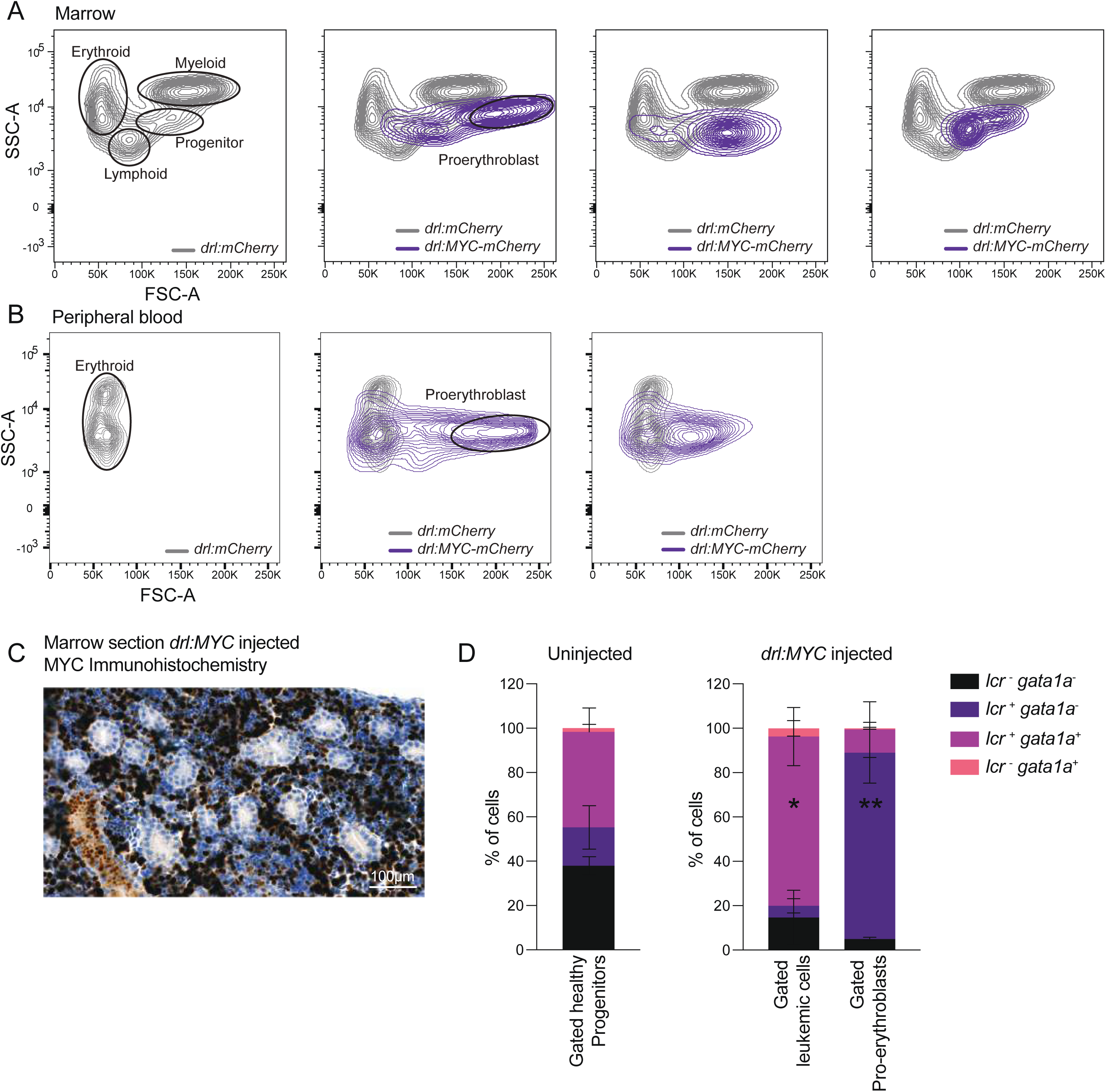
MYC overexpression induces expansion of erythroid progenitors. (A) Flow cytometry contour plot of a representative example of *drl:mCherry* control (grey) overlapped with a three examples of *drl:MYC:mCherry* leukemic (purple) marrow showing leukemic cell and pro-erythroblast gating example and emphasizing the mosaicism of our system. (B) Flow cytometry contour plot of a representative example of *drl:mCherry* control (grey) overlapped with a two examples of *drl:MYC:mCherry* leukemic (purple) peripheral blood samples showing circulating leukemic cell and pro-erythroblast. (C) MYC immunohistochemistry on a *drl:MYC* marrow section showing a packed marrow with MYC^+^ leukemic cells. (D) Bar graph quantifying the mean with standard devation of the fraction of *lcr:GFP* and *gata1a:dsRed* populations in healthy progenitors (left panel, n=4), leukemic cells and proerythroblasts (right panel, n=3) demonstrating the erythroid identity of transformed populations. * p<0.05 between healthy *vs*. *drl:MYC lcr^+^ gata1a^+^* and ** p<0.001 between healthy *vs*. *drl:MYC lcr*^+^ *gata1a*^-^.

**Fig. S2.**
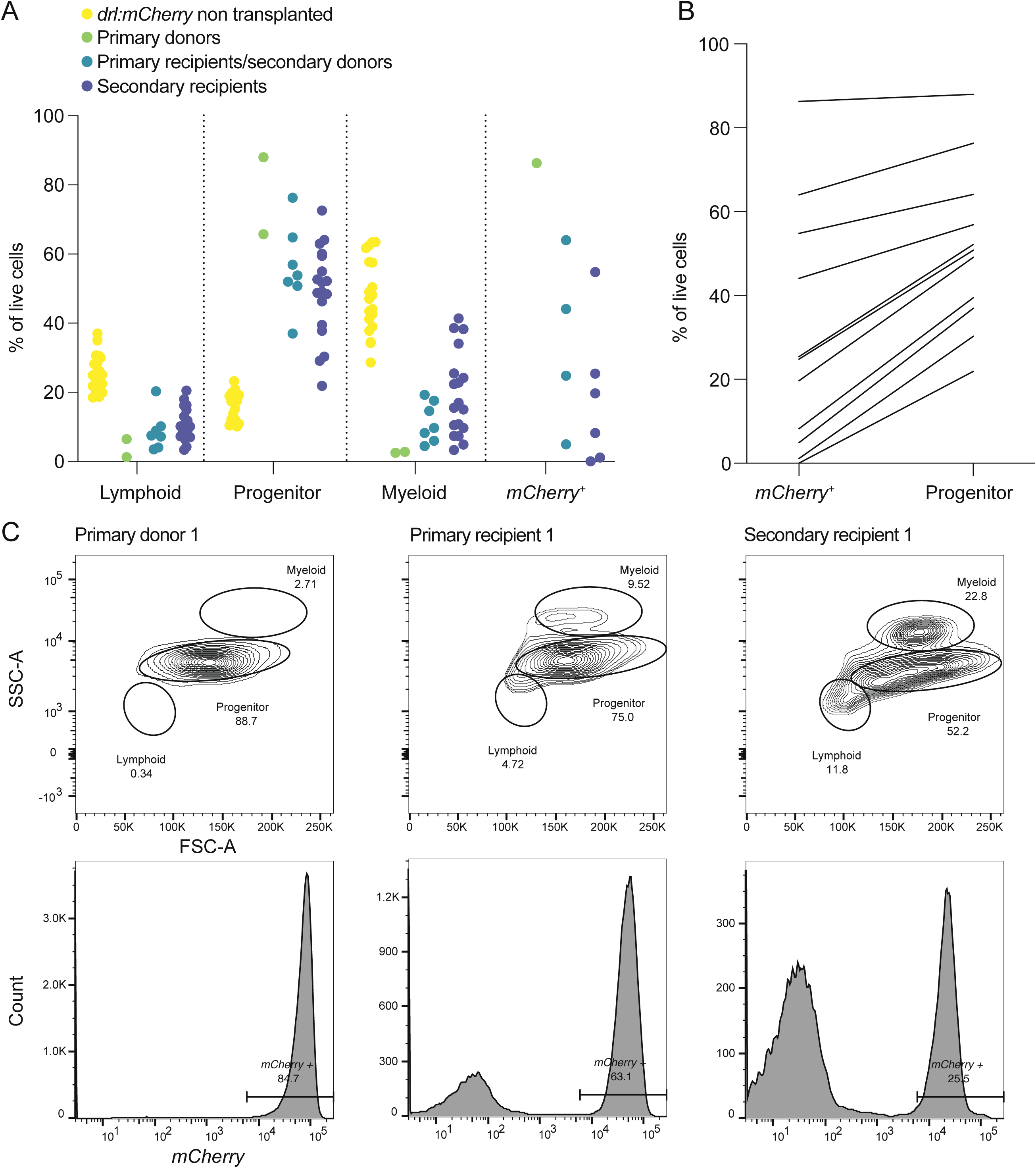
*drl:MYC* leukemic cells are serially transplantable. (A) Scatter plot quantifying the percent of live cells in lymphoid, progenitor, myeloid gates as defined by FSC/SSC and mCherry gate in non-transplanted controls (n=19), primary donors (n=2), primary recipients (n=7) and secondary recipients (n=18). (B) Line graph demonstrating the correlation between *mCherry^+^* cells and cells overlapping with the progenitor gate as defined in FSC/SSC space demonstrating that both readouts can be used to measure engraftment. (C) Flow cytometry plots of a representative example of donor 1 (left panels) and a subsequent engrafted primary recipient (middle panels) and secondary recipient (right panels). Top row shows contour plots of live cells in FSC/SSC space and percent in lymphoid, myeloid and progenitor gate are indicated under each gate. Bottom row shows histograms of live cells gated for *mCherry* and percent of *mCherry^+^*cells are indicated above each gate.

**Fig. S3.**
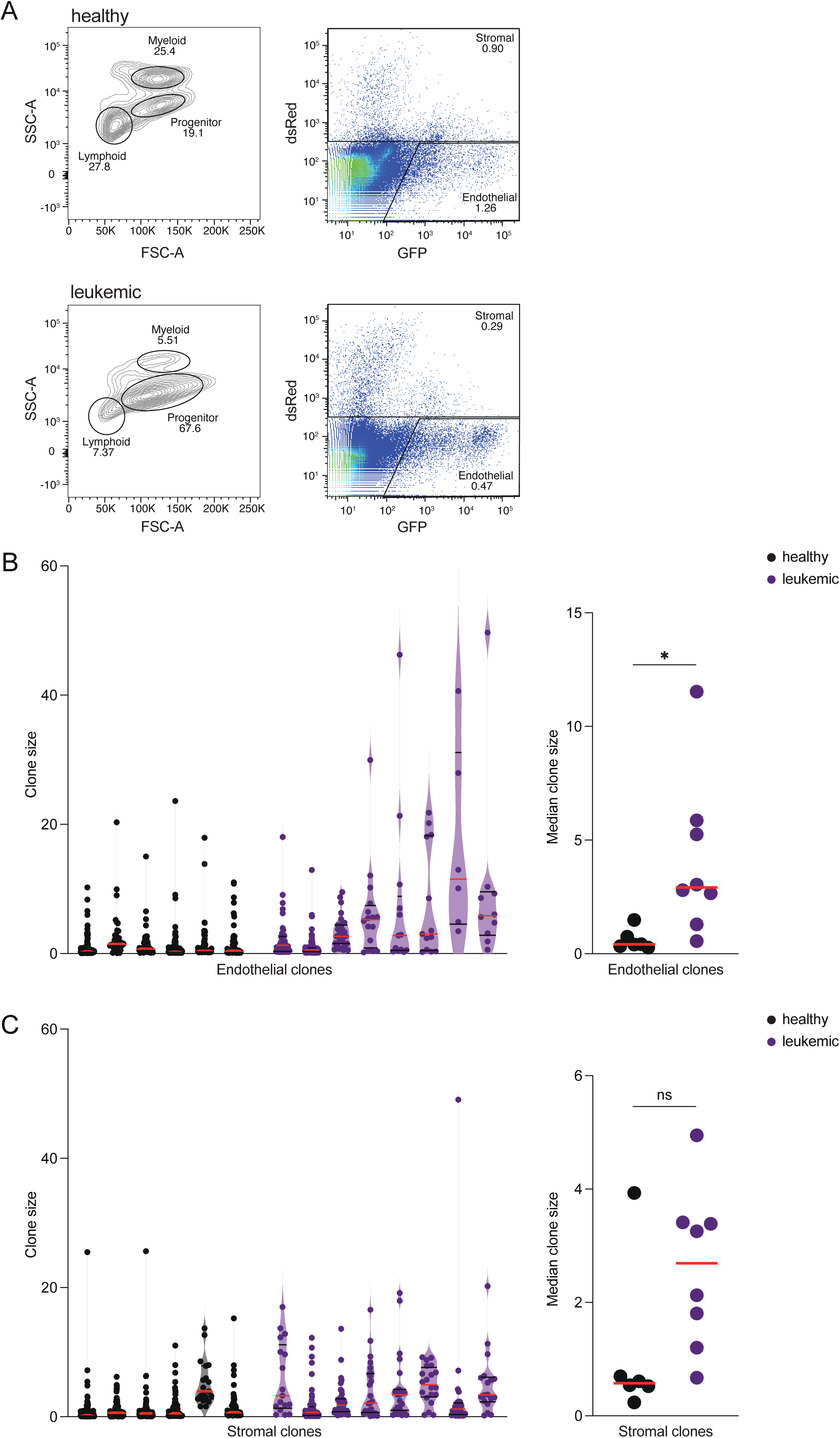
ECs and MSCs from leukemic marrows are clonally expanded. (A) Flow cytometry plots of a representative example of healthy (top panels) and leukemic (bottom panels) enzymatically dissociated marrows in FSC/SSC space (contour plots in left panels) and *GFP* and *dsRed* space (pseudo-color plots in right panels). Under each gate is indicated the percent of live cells. For GESTALT analysis, cells from endothelial and stromal gates were sorted. (B) Violin plot of EC clone size of individual healthy (n=6) and leukemic (n=8) marrows revealing clonal selection upon leukemia (left panel) and median EC clone size for healthy and leukemic fish (right panel). Red line indicates median. * p<0.05 (C) Violin plot of MSC clone size of individual healthy (n=6) and leukemic (n=8) marrows revealing clonal selection upon leukemia (left panel) and median MSC clone size for healthy and leukemic fish (right panel). Red line indicates median. ns: non-significant.

**Fig. S4.**
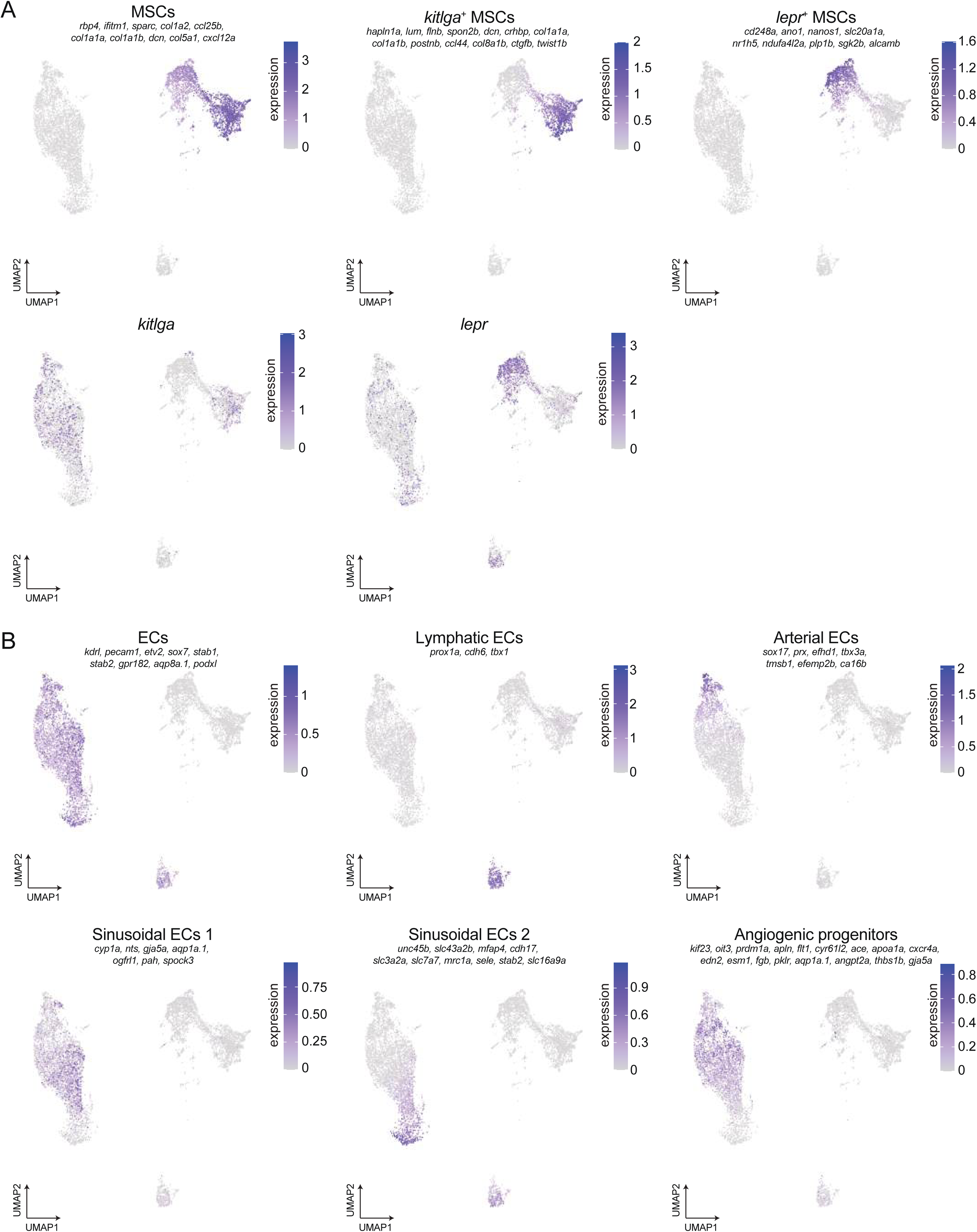
Distinct MSC and EC subpopulations can be identified by scRNA-seq. (A) UMAP dimensionality reduction (n=9,176 cells) as defined in Fig. 2E. showing the expression of the MSC gene signature (top left), the *kitlga*^+^ MSCs signature (top middle), the *lepr*^+^ MSC signature (top right), *kitlga* (bottom left) and *lepr* (bottom middle) in healthy and leukemic marrows combined on the same UMAP. Gene signatures are listed under the sub-population name. Single genes are indicated above their respective plots. Color represents signature or gene expression level. (B) UMAP dimensionality reduction (n=9,176 cells) as defined in Fig. 2E. showing the expression of the EC gene signature (top left), the lymphatic EC signature (top middle), the arterial EC signature (top right), the sinusoidal EC 1 signature (bottom left), the sinusoidal EC 2 signature (bottom middle) and the angiogenic signature (bottom right) in healthy and leukemic marrows combined on the same UMAP. Gene signatures are listed under the sub-population name. Color represents signature or gene expression level.

**Fig. S5.**
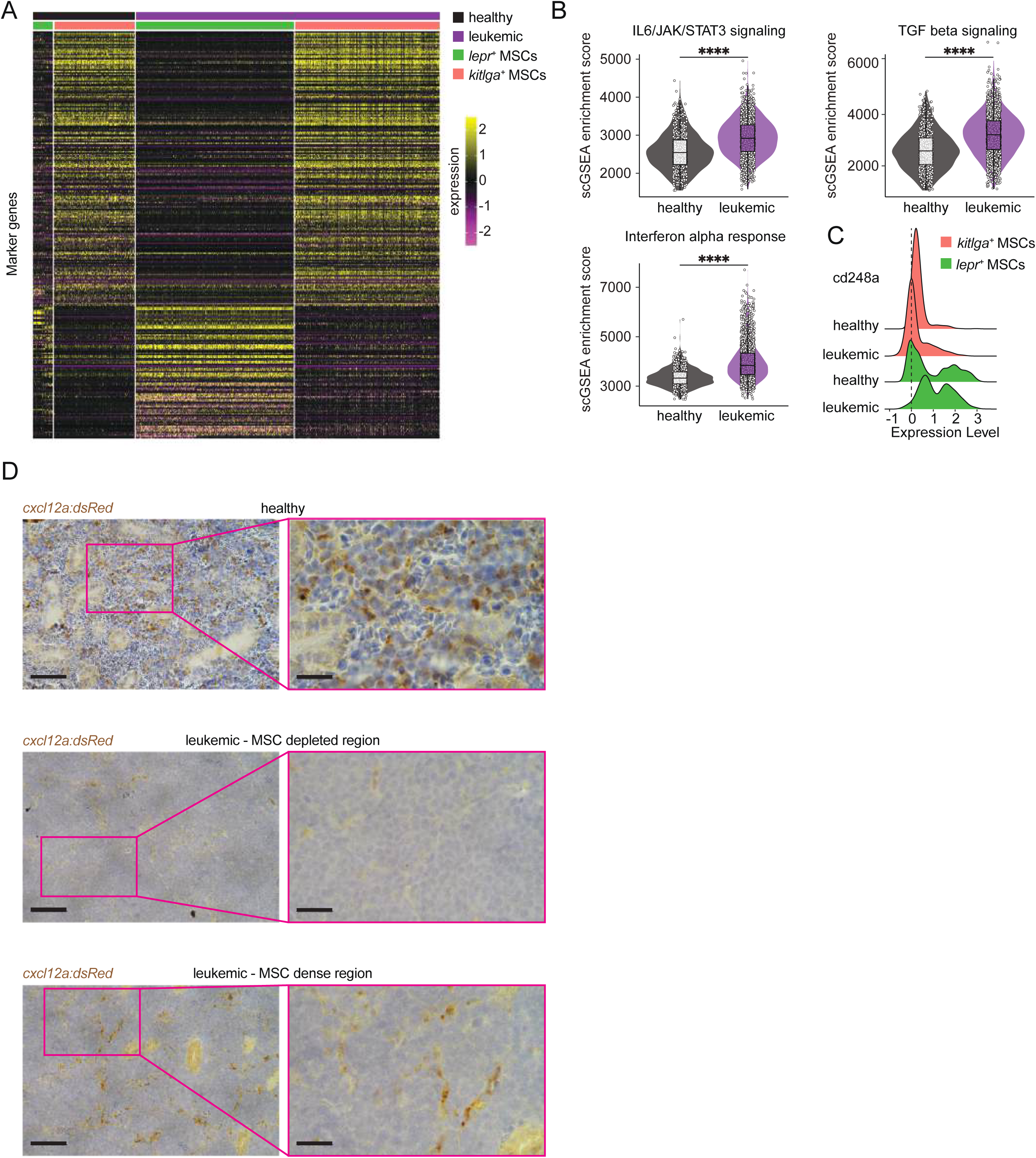
*lepr*^+^ MSCs are activated in leukemic marrows. (A) Heatmap of differentially expressed genes (p-value < 0.05) in *lepr*^+^ and *kitlga*^+^ MSC sub-populations in healthy and leukemic marrows. *kitlga*^+^ MSCs gene expression is unchanged upon leukemia while *lepr*^+^ MSCs undergo a transcriptional change (bottom blocks on heatmap). Heatmap color represents gene expression level. Top color bar represents MSC subpopulation and sample identity. (B) Violin plots of scGSEA pathways identified as differentially enriched in healthy versus leukemic MSCs and involved in MSC function and activation. **** p<0.0001. (C) Stacked histogram of *cd248a* activation marker gene expression in *lepr*^+^ and *kitlga*^+^ MSC sub-populations in healthy and leukemic marrows revealing specific and upregulated expression in *lepr*^+^ MSCs. (D) Representative example of immunohistochemistry for dsRed fluorescent protein on sections from healthy and leukemic *cxcl12a:dsRed* transgenic zebrafish showing distinct regions of MSC density. Scale bar 100μm (left panels) and 20μm (right panels).

**Fig. S6.**
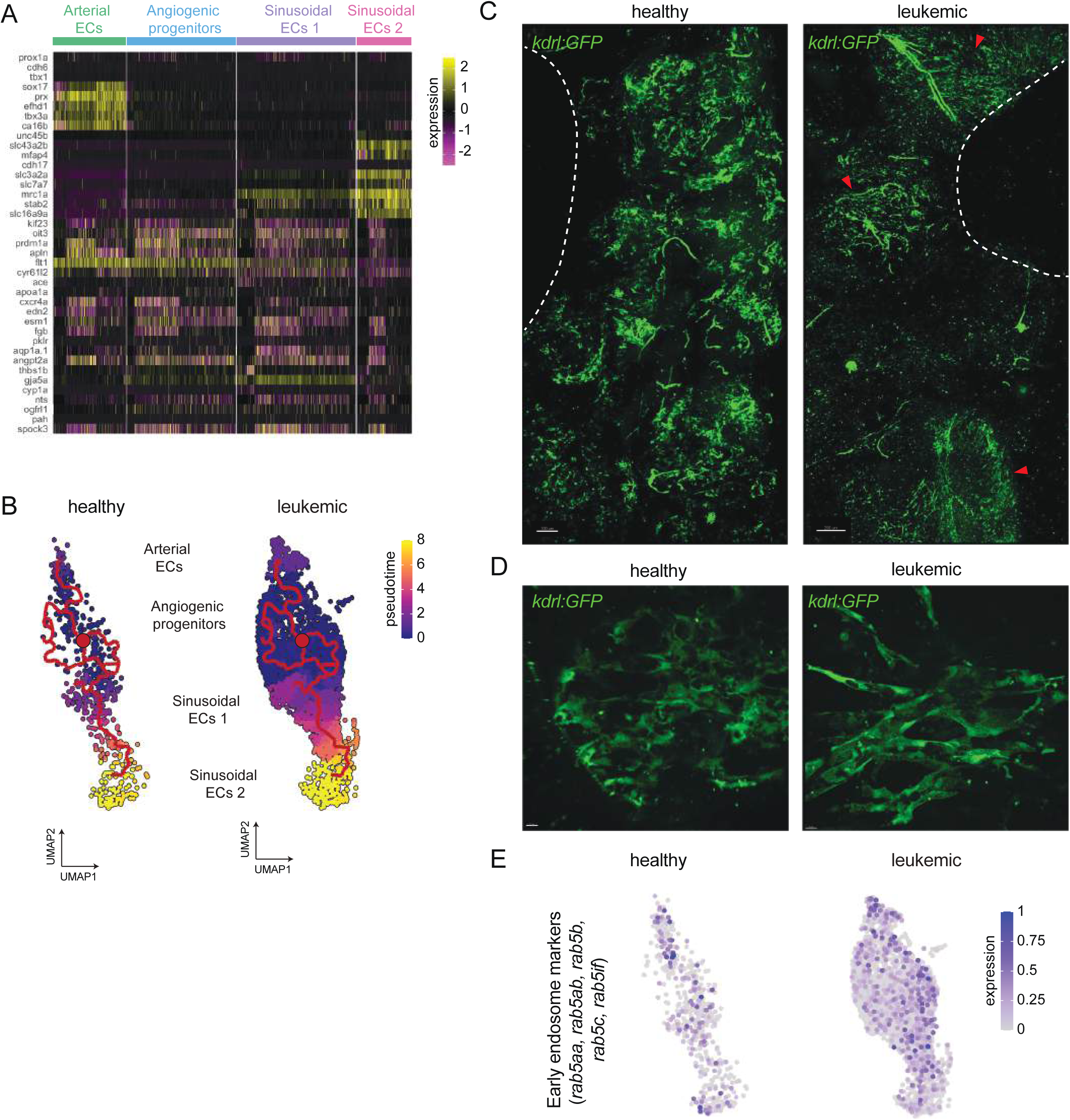
Leukemia induces expansion of angiogenic progenitors. (A) Heatmap of the top 8 marker genes in arterial ECs, angiogenic progenitors, sinusoidal EC 1 and 2 subpopulations demonstrating that angiogenic progenitors share marker genes with all other populations. Heatmap color represents gene expression level. Top color bar represents EC subpopulations. (B) UMAP of *in silico* purified ECs after Monocle 3 trajectory analysis starting at the angiogenic progenitor cluster demonstrating two distinct differentiation paths towards arterial or sinusoidal ECs. Red line is monocle inferred trajectory and UMAP is colored based on pseudotime. (C) Representative image of live marrow from healthy and leukemic *kdrl:GFP* transgenic fish demonstrating a change in vasculature structure with dense and depleted regions. White line isolate regions without tissue. Red arrows indicate vascular dense regions. Scale bar 100μm (left panel) and 200μm (right panel) (D) Representative images of close-ups (100X) from dense vascular regions of healthy and leukemic marrows demonstrating a change in structure and endocytic content of ECs. Scale bar 10μm (both panels) (E) UMAP dimensionality reduction as defined in Fig. 2E. showing the expression of early endosome markers demonstrating upregulation in leukemic ECs. Color represents combined gene expression level.

**Fig. S7.**
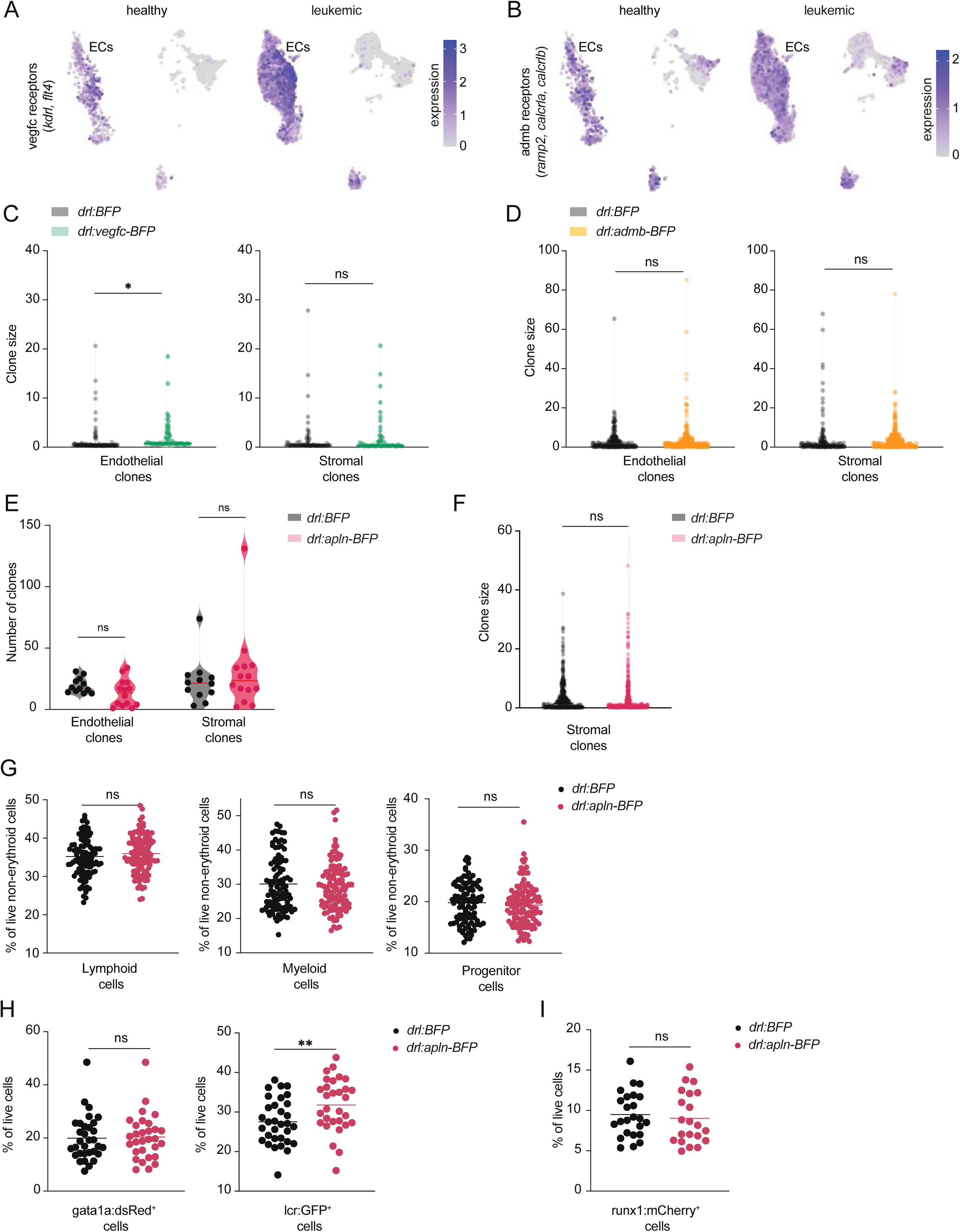
*apln* overexpression induces EC clonal selection and alters HSPC clonal output. (A) UMAP dimensionality reduction (n=9,176 cells) as defined in Fig. 2E. showing the expression of the two *vegfc* receptors *kdrl* and *flt4* in healthy and leukemic marrows demonstrating selective and increased expression in leukemic ECs. Color represents combined gene expression level. (B) UMAP dimensionality reduction (n=9,176 cells) as defined in Fig. 2E. showing the expression of the three *admb* receptors *ramp2*, *calcrla* and *calcrlb* in healthy and leukemic marrows demonstrating increased expression in leukemic ECs. Color represents combined gene expression level. (C) Violin plot of EC (left panel) and MSC (right panel) clone size of pooled control *drl:BFP* (n=6) and *drl:vegfc-BFP* (n=6) marrows revealing EC clonal expansion driven by relatively small clones. * p<0.05. ns=non-significant. (D) Violin plot of EC (left panel) and MSC (right panel) clone size of pooled control *drl:BFP* (n=5) and *drl:admb-BFP* (n=11) marrows revealing no EC or MSC clonal expansion. ns=non-significant. (E) Violin plot of the number of clones detected in purified ECs (left panel) and MSCs (right panel) in *drl:BFP* control (n=12) and *drl:apln-BFP* marrows (n=14) showing no significant decrease in clone number. The red line indicates the median value. ns= non-significant. (F) Violin plot of MSC clone size of pooled control *drl:BFP* (n=12) and *drl:apln-BFP* (n=14) marrows revealing no MSC clonal expansion. ns=non-significant. (G) Dot plot of live marrow cells quantified by flow cytometry and gated into three distinct populations demonstrating no changes in hematopoietic lineages in *drl:apln-BFP* marrows (n=115) compared to *drl:BFP* controls (n=102). Black line shows mean. ns=non-significant. (H) Dot plot of live marrow cells quantified by flow cytometry and gated for *gata1a:dsRed* erythroid progenitors and *lcr:GFP* erythrocytes showing a significant increase in erythrocytes in *drl:apln-BFP* marrows (n=30) compared to *drl:BFP* controls (n=32). Black line shows mean. **p<0.01. ns=non-significant. (I) Dot plot of live marrow cells quantified by flow cytometry and gated for *runx1+23:mCherry* HSPCs showing no difference in *drl:apln-BFP* marrows (n=21) compared to *drl:BFP* (n=24) controls. Black line shows mean. ns=non-significant.

**Fig. S8.**
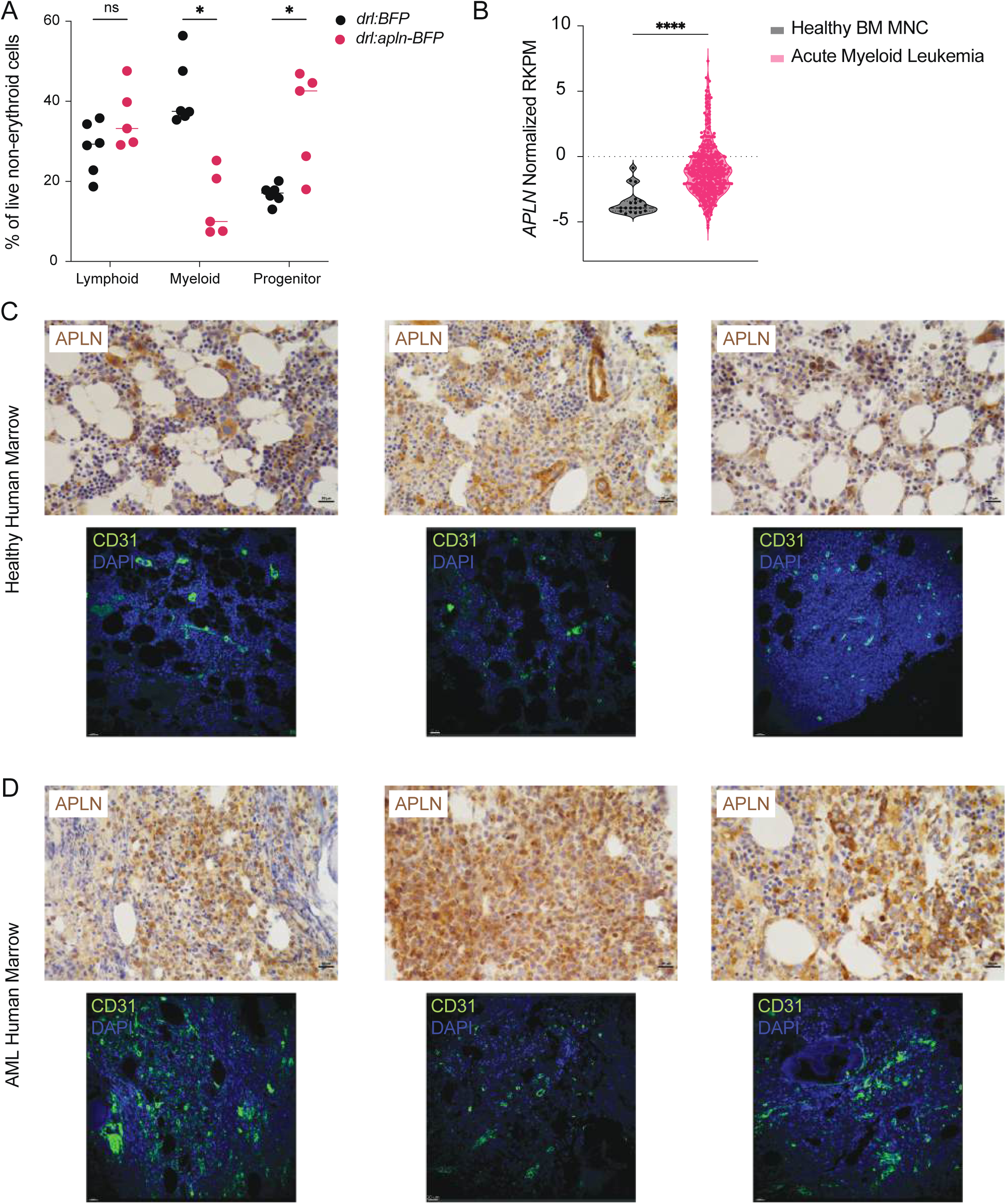
Human AML express high levels of APLN and have expanded vasculature. (A) Dot plot of live marrow cells quantified by flow cytometry and gated into three distinct populations demonstrating changes in hematopoietic lineages in abnormal *drl:apln-BFP* marrows (n=5) compared to *drl:BFP* controls (n=5). * p<0.05. ns=non-significant. (B) Violin plot of *APLN* normalized RPKM from the BEAT AML study from healthy bone marrow mononucleated cells (n=19) and AML cells (n=440). **** p<0.0001. (C) APLN immunohistochemistry (top panels) and CD31 immunohistochemistry (bottom panels) of healthy human marrows. Scale bar 20μm (all panels). (D) APLN immunohistochemistry (top panels) and CD31 immunohistochemistry (bottom panels) of AML human marrow. Scale bar 20μm (all panels).

**Supplementary table 1. Marker genes for EC and MSC subpopulations.** Genes included are significantly upregulated in each population (p<0.05).

